# A collection of *Serratia marcescens* differing in their insect pathogenicity towards *Manduca sexta* larvae

**DOI:** 10.1101/2020.07.29.226613

**Authors:** Ellen C. Jensen, Diana Sue Katz Amburn, Aimee Hultine Schlegel, Kenneth W. Nickerson

## Abstract

We investigated the ability of *Serratia marcescens* to kill *Manduca sexta* (tobacco/tomato hornworm) larvae following injection of ca. 5 × 10^5^ bacteria into the insect hemolymph. Fifteen bacterial strains were examined, including 12 non-pigmented clinical isolates from humans. They fell into 6 groups depending on the timing and rate at which they caused larval death. Relative insect toxicity was not correlated with pigmentation, colony morphology, biotype, motility, capsule formation, iron availability, surfactant production, swarming ability, antibiotic resistance, bacteriophage susceptibility, salt tolerance, nitrogen utilitization patterns, or the production of 4 exoenzymes: proteases, DNase, lipase, or phospholipase. There were marked differences in chitinase production, the types of homoserine lactone (HSL) quorum sensing molecules produced, and the blood agar hemolysis patterns observed. However, none of these differences correlated with the six insect larval virulence groups. Thus, the actual offensive or defensive virulence factors possessed by these strains remain unidentified. The availability of this set of *S. marcescens* strains, covering the full range from highly virulent to non-virulent, should permit future genomic comparisons to identify the precise mechanisms of larval toxicity.

## Introduction

*Serratia marcescens* has been of interest to microbiologists for many years. Much of this interest derives from the production of a bright red, blood-like pigment called prodigiosin by many strains [1]. This red pigmentation led to countless deaths from “bleeding host” hysteria during the Middle Ages [2] as well as providing the name for “red diaper syndrome” [3]. In early studies on the feasibility of bacteriological warfare, this distinctive pigmentation was the rationale for *S. marcescens* being chosen as the test organism for aerial spraying over populated areas of the United States [4]. *S. marcescens* is estimated to cause 2.3% of the nosocomial infections in the United States [5].

However, *S. marcescens* is also a well known insect pathogen [6, 7] which has been reported to cause disease in at least 100 species of insects [8]. The insect pathogenicity of *S. marcescens* has a renewed urgency in terms of the global honey bee colony decline [9, 10]. Raymann et al [10] suggested that *S. marcescens* is a widespread opportunistic pathogen of adult honey bees and that the bees are likely more susceptible following perturbation of their normal gut microbiota, the presence of Varroa mites, and exposure to various antibiotics or pesticides [10]. We are interested in the biochemical and physiological features contributing to insect virulence and we used *S. marcescens* in our study showing the involvement of eicosanoids in the response of *Manduca sexta* larvae to bacterial infection [11]. Eicosanoids are signaling metabolites derived from C_20_ polyunsaturated fatty acids. In this work we injected *S. marcescens* cultures into the larval hemolymph at ca 5 × 10^5^ bacteria per larva and then followed mortality as well as the ability or inability of larvae to clear the pathogenic bacteria from their hemolymph. We observed that dexamethasone, an inhibitor of phospholipase A_2_, significantly reduced the ability of larvae to clear pathogenic bacteria while this ability was restored by treatment with arachidonic acid, the C_20:4_ fatty acid released by phospholipase A_2_. We concluded that eicosanoids likely mediate invertebrate immune responses [11].

Naturally, we were concerned when another laboratory told us that they were unable to repeat our findings. These difficulties were traced to the strains of *S. marcescens* being used. When they used our strain (now called KWN) they could repeat our findings. This was the first indication we had that the strain of *S. marcescens* used in our eicosanoid studies [11] was unusually pathogenic to insects. This realization led us to compare 15 isolates of *S. marcescens* with regard to their pathogenicity towards *M. sexta* larvae. The bacteria formed 6 pathogenicity groups, ranging from highly pathogenic to non-pathogenic. Strain KWN was in the highly pathogenic group. We now report how these isolates differ from one another while providing hints regarding the biochemical and physiological factors responsible for these widely differing pathogenicities. Although the precise mechanisms of their pathogenicity differences have not yet been identified, the collection should be of use because its members cover stepwise gradations from being highly pathogenic to non-pathogenic.

## Materials and Methods

### Organisims and Cultural Conditions

For *Serratia marcescens*, strain KWN was isolated from the Teresa Street Sewage Plant in Lincoln, NE by Drs. Ellen Jensen and Bruce Lahm in 1986; strain D1 was from Presque Isle Cultures, Erie, PA; and strain Nima was obtained (1976) from the late Prof. R.P. Williams, University of Texas, Austin, TX. D1 is a mutant derived from Nima. All the rest were clinical isolates obtained from three hospitals in Kansas [12]. The bacteria were grown in Luria-Bertani broth (Difco Detroit) at 30-32°C with rotary agitation at 120 rpm. Bacteria used in insect bioassays were always in exponential phase growth. For the TLC assay for autoinducers [13], the reporter strains were *Chromobacterium violaceum* CV026 from Drs. Yan Jiang and Paul Williams, University of Nottingham, *Agrobacterium tumefaciens* NT1 (pDCI 41E33) from Dr. Stephen Farrand, University of Illinois, and *E. coli* MG4 (pKDT17) from Dr. Barbara Iglewski, University of Rochester.

### Insect Bioassay

Eggs of *Manduca sexta*, the tobacco hornworm, were obtained from the USDA, Fargo, ND (pre 1995), the USDA, Beltsville, MD (1995 and 1996), and Carolina Biological Supply, Burlington, NC (1997). Larvae were reared under standard conditions at 28°C with a 16 hr light/8 hr dark photoperiod [14]. Each strain of *S. marcescens* was bioassayed in triplicate experiments with ten early fifth/last instar prewandering larvae. The larvae were injected as described [11] with 10 µl suspensions (ca. 5 × 10^5^ bacteria) and then examined at 2 hour intervals to see if they could respond to stimuli. The larval response to strain KWN was the same for all three sources of *M. sexta* (14 experiments over a 7 year period) and, consequently, we do not believe that the genetic background of the insect host influences our results.

### Capsule production

Cultures were grown in Gauger’s G medium [15], a defined glucose-salts medium consisting of: 20g glucose, 2g asparagine, 0.5g KH_2_ PO_4_, and 0.28g Mg SO_4_ per L of distilled water, adjusted to pH 6.8, for 3 days at 30°C. This medium has a high C/N ratio and promotes excellent capsule formation in *Enterobacter cloacae* [16]. The negative staining procedure outlined by Chan et al [17] was used. A loopful of culture was mixed with nigrosin, spread on the slide, dried, fixed with methanol, counterstained with crystal violet, rinsed gently, air dried, and observed under oil immersion (1000x). The relative capsule diameter was determined microscopically using an uncalibrated ocular micrometer scale.

### Motility, swarming, wettability, and serrawettin

Motility and swarming abilities were determined as described by Alberti and Harshey [18] while wettability and biosurfactant measurements followed Matsuyama et al [19]. The motility and swarming studies were done on LB agar plates containing 0.35 and 0.75% agar, respectively.

### Production of Extracellular Enzymes

We screened for five extracellular enzymes: chitinases, DNases, proteases, lipases, and phospholipases. Each assay measured the diameter of the zone of clearing formed around colonies on solid media. The chitin-containing plates for the detection of chitinase were prepared as described by Sundheim et al [20], the skim milk and gelatin-containing plates for the detection of proteases were as described by Kelley and Post [21], and the DNase plates with toluidine blue were as described by Chen et al [22]. The lipase plates were prepared as described by Lovell and Bibel [23]: 1% peptone, 0.5% NaCl, 0.01% CaCl_2_ - 2H_2_O, and 1.6% agar, adjusted to pH 7.4, autoclaved, and cooled to 50°C, whereupon 1 ml of Tween which had been autoclaved separately and cooled to 50°C was added per 100 ml. Separate lipase plates were prepared with Tweens 20, 40, 60, and 80. Finally, the phosphatidyl choline-containing plates for the detection of phospholipases were of our own design. Nutrient agar (Difco, Detroit) was suspended in 85% of the usual volume of water, autoclaved, and cooled to 50°C. The phosphatidyl choline was prepared as a 2% solution in ethanol, diluted with 2 volumes of sterile water preheated to 50°C, and mixed with the molten agar at a ratio of 15:85. In all cases the test cultures were grown overnight in nutrient broth (Difco, Detroit) and inoculated as a single spot on a 90mm Petri dish using a sterile applicator stick. Six cultures were tested per plate. The plates were incubated at 30°C and observed after 24, 48, and 72 hours. In many cases duplicate series of plates were incubated in both air (regular) and candle jar environments on the assumption that the high CO_2_/low O_2_ conditions in the candle jar better simulated those in the insect hemolymph.

### Nitrogen Source

Bacteria were grown in Bacto Yeast Carbon Base (Difco, Detroit) supplemented with 10 mM ammonium sulfate, potassium nitrate, urea, uric acid, or allantoin, and adjusted to pH 7.0 prior to use.

### Biotyping

Biotyping based on carbon source utilization was done by the method of Grimont and Grimont [24]. The strains were compared with regard to their ability to use benzoic acid, D,L-carnitine, meso-erythritol, 3-hydroxybenzoic acid, 4-hydroxybenzoic acid, and trigonelline as their sole source of carbon and energy.

### Autoinducer Production

The strains of *S. marcescens* tested for autoinducer production were grown with shaking at 30°C in either Luria broth, Trypticase soy broth, or CCY medium. CCY is a nutrient limiting, modified sporulation medium [25]. The cultures were grown to stationary phase and pelleted by centrifugation (8,000 rpm for 20 min). The supernatants were filtered through 0.45µm nitrocellulose filters and extracted into an equal volume of ethyl acetate containing 0.04% glacial acetic acid. Portions (1.5ml) of the ethyl acetate were transferred to microcentrifuge tubes and the ethyl acetate was removed in a Savant SVC100H Speed Vac Concentrator. The residues were resuspended in 20 ul of ethyl acetate whereupon samples (4ul) were applied to C_18_ reversed-phase TLC plates (Baker 7013) and the chromatograms were developed with methanol/water (60:40 v/v) [13]. The three reporter strains were *C. violaceum* CV026, *A. tumefaciens* NT1, and *E. coli* MG4. This *E*.*coli* is a reporter strain for the C_12_PAI 1 from *Pseudomonas aeruginosa* [26]. After development, the dried TLC plates were overlaid with a culture of the reporter bacterium in 0.75% agar [47]. The *A. tumefaciens* and *E. coli* agar overlays contained 60ug/ml of X-Gal [13]. The plates were incubated at 28°C (*C. violaceum* CV026 and *A. tumefaciens* NT1) or 37°C (*E. coli* MG4) in closed plastic containers and observed for color development at 6 hr intervals. The samples applied (4ul) are equivalent to 12.5 ml of culture for the *S. marcescens* strains and 10 ml of culture for the positive controls.

### Antibiotic sensitivity/resistance

The methods employed followed those we used previously to characterize antibiotic resistance in bacteria isolated from larval guts of the oil fly *Helaeomyia petrolei* [27]. The discs contained: 10 µg ampicillin, 100 µg piperacillin, 30 µg cefoxitin, 30 µg cefotaxime, 30 µg aztreonam, 10 µg imipenem, 10 IU bacitracin, 30 µg vancomycin, 300 IU polymyxin B, 10 µg colistin, 25 µg sulfamethoxazole/trimethoprim, 10 µg streptomycin, 30 µg kanamycin, 30 µg neomycin, 10 µg tobramycin, 30 µg tetracycline, 30 µg chloramphenicol, 15 µg erythromycin, 5 µg rifampin, 30 µg nalidixic acid, 5 µg ciprofloxacin, 10 µg norfloxacin, 300 µg nitrofurantoin, or 5 µg novobiocin. The imipenem and sulfamethoxazole-containing discs were from Oxoid, Ltd, Basingstoke, UK. All the rest were Sensi-discs from Becton Dickinson/BBL, Sparks, MD.

### Bacteriophage sensitivity/resistance

The seven broad host range bacteriophage described by Jensen et al [28] (SN-1, SN-2, SN-X, SN-T, BHR1, BHR2, and D_3_C_3_) were grown in both *Sphaerotilus natans* and *Pseudomonas aeruginosa*, whereupon all 15 strains of *S. marcescens* were challenged for both plaque formation (agar plates) and increased bacteriophage titer (liquid). All procedures followed those described by Jensen et al [28].

## Results

### Insect Bioassays

The 15 strains of *S. marcescens* tested fell into six groups with regard to their pathogenicity to *M. sexta* larvae (Fig 1 and Table 1). Strain D1 was non-toxic, strains KWN and 9674 were highly toxic, and the other 12 strains showed intermediate levels of toxicity (Table 1). With constant numbers of bacteria introduced, the 6 pathogenicity groups differed in the times at which larval death was first observed and the subsequent rates of larval death (Fig 1). After injection, the infected larvae stopped feeding within one hour, stopped moving, and became flaccid. No signs of hemolymph bleeding from the injection sites were observed [29]. After death, if the infecting *S. marcescens* was pigmented, the insect cadavers also acquired a dark red pigmentation. For those strains of *S. marcescens* which did not give 100% killing (Group 5 in Fig 1), all of the insects stopped feeding within one hour and then, 8-12 hours later, some of them started feeding again and became survivors. Only one of the clinical isolates, strain 9674, joined strain KWN in the highly virulent category (Table 1). Strain KWN was toxic by injection but not when incorporated into the larval diet or when painted on the outer surface of the larvae. This distinction is consistent with the lack of chitinase production by strain KWN (Table 1). For strain KWN, we found no change in toxicity using three different sources of *M. sexta* eggs over a period of seven years, or if the bacteria were in exponential phase or stationary phase at the time of injection, or if they were injected in 10ul LB broth (standard) or in 10 µl of 1 mM EDTA. Similarly, strain D1 remained non-toxic when injected in 10 µl of 1-100 µM FeSO_4_ 7H_2_O. These latter additions were designed to test whether, as in vertebrates [30], bacterial pathogenicity is correlated with the availability of iron. No correlation between pathogenicity and iron availability was observed.

**Table 1.** Strains of *Serratia marcescens* used in this study.

| Strain | Insect<br>Pathogenicity<sup>a</sup> | Biotype<sup>b</sup> | Pigmented | Chitinase<br>Production | Motility | Swarming | Capsule |
| --- | --- | --- | --- | --- | --- | --- | --- |
| KWN | 1 | A1a | + | - | + | + | 1.5 |
| 9674 | 1 | A8b | - | - | + | + | 1.5 |
| E223 | 2 | TCT | - | + | + | +<sup>vs</sup> | 1 |
| 1682 | 2 | A5 | - | + | + | + | 2.5 |
| 1-2232b | 2 | A4a | - | - | + | +<sup>s</sup> | 1 |
| 661 Ulrich | 2 | A4a | - | - | + | + | 1 |
| 00672A | 3 | A4a | - | + | + | + | 2 |
| 5384 | 3 | A8a | - | + | + | + | 1.5 |
| D163 | 3 | A3a | - | + | + | + | 1.5 |
| Figus | 3 | A4a | - | + | + | + | 2 |
| 968A | 4 | A4a | - | + | + | + | 1.5 |
| 131 Watkins | 4 | A8a | - | + | + | - | 2 |
| 2698B | 5 | A3b | - | + | + | + | 1.5 |
| NIMA | 5 | A2a | + | + | + | - | 1.5 |
| D1 | 6 | A2a | + | + | - | - | 1 |

a/ groups arranged with group 1 being the most pathogenic and group 6 being non-pathogenic
b/ as described by Grimont and Grimont [24]
+<sup>s</sup> = positive but slow; swarming required 30 hours
+<sup>vs</sup> = positive but very slow;swarming required 48-72 hour

**Fig 1.**
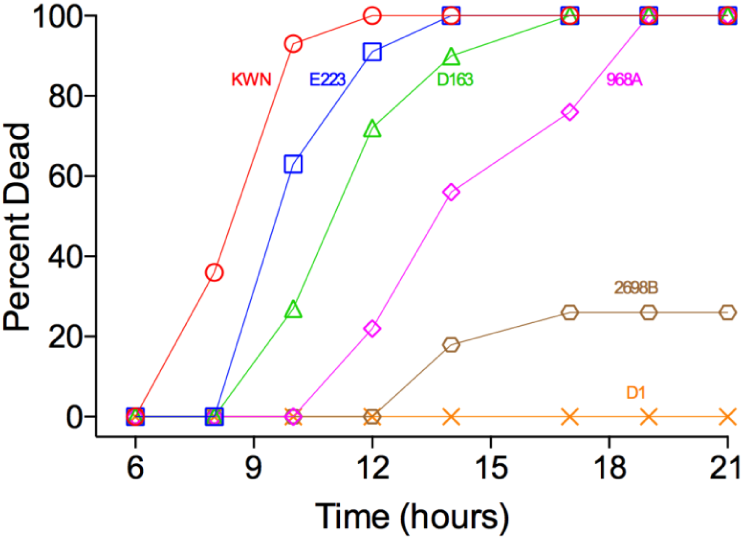
Kinetics of *Manduca sexta* larval death following injection of *Serratia marcescens*. Each larva was injected with ca. 5 × 10^5^ bacteria. Symbols: **○** = strain KWN; □ = strain E223; Δ = strain D163; ◊ = strain 968A; open hexagon = strain 2698B; and X = strain D1. Fifteen strains of *S. marcescens* were tested (triplicate assays of 10 larvae each -- total 30 larvae). These 6 strains were chosen to illustrate the 6 different rates of larval death listed in Table 1.

### Pigments, Capsules, Motility, and Surfactants

We next sought to determine what features of strains KWN and 9674 made them more toxic than the other strains of *S. marcescens*. There were no obvious differences in colony morphology among the strains except for those that were pigmented red (Table 1). However, the red pigment prodigiosin was not a virulence factor towards *M. sexta* larvae in that the pigmented strains included both the most toxic and least toxic strains (Table 1). This conclusion agrees with the observations of Zhou et al [31] who found that 8 pigmented and non-pigmented strains of *S. marcescens* had equivalent LD_50_ values towards larvae of the silkworm *Bombyx mori*. In agreement with the findings of Grimont and Grimont [24, 32], the red pigmentation by prodigiosin was only observed for biotypes A1 and A2 (Table 1).

All strains produced large capsules when grown on the defined, high C/N Gauger’s medium [16]. Capsules of roughly equivalent size were observed by negative staining [17] for all strains (Table 1). We also compared the strains with regard to their motility on semi-solid 0.35% agar plates and their swarming ability on 0.75% agar plates [18]. All strains were motile except for the non-toxic strain D1 and, while most of the strains were capable of swarming, there did not appear to be a correlation between swarming and insect pathogenicity (Table 1). Many swarming bacteria synthesize and secrete surfactants which enable those bacteria to spread over surfaces [19, 33,34]. For *Serratia sp*. this surfactant is often the lipopeptide serrawettin [19, 33, 34] and thus we examined our 15 strains of *S. marcescens* for serrawettin production and wettability on four surfaces (S1 Table). Individual strains produced each of the 3 serrawettins (W1, W2, and W3) identified by thin-layer chromatography by Matsuyama et al [19] while strain D163 produced a unique TLC spot which we have called W4 (S1Table). However, there was no correlation between serrawettin production and insect pathogenicity. For instance, the highly pathogenic strain E223 had no detectable serrawettin or wettability (S1Table) and only late developing swarming (Table 1).

### Hemolysis and Salt Tolerance

Many bacterial pathogens release hemolysins to lyse erythrocytes and other cell types so that they gain access to the nutrients in those cells, especially iron and hemoglobin. When our 15 strains of *S. marcescens* were examined on blood agar plates (Table 2), three exhibited α-hemolysis, six exhibited β-hemolysis (complete zones of clearing), and one (Figus) exhibited double hemolysis with both α- and β-rings, and five exhibited no hemolysis, which paradoxically is called γ-hemolysis. However, there was no correlation between larval virulence and hemolysis. In particular, both the most virulent strains (KWN and 9674) and the least virulent strains (2698B and D1) did not exhibit hemolysis (Table 2). Grimont and Grimont [24] found the same four hemolysis patterns. In their analysis of 2, 210 isolates of *S. marcescens* they noted that β-hemolysis was strongly associated with biotype A4 (80% of A4 strains as opposed to 0.7% of strains of other biotypes). This generalization also held for our work in that four of the six β-hemolytic strains were biotype A4a as was the doubly hemolytic Figus (Table 2).

**Table 2.** Blood agar hemolysis and salt tolerance of *Serratia marcescens*.

| Strain | Hemolysis <sup>a/</sup> | Growth on Mannitol-Salt Agar |  |  |
| --- | --- | --- | --- | --- |
|  |  | 25°C | 30°C | 37°C |
| KWN | gamma | ++ | + | - |
| 9674 | gamma | ++ |  |  |
| E223 | beta | ++ | +/- | +/- |
| 1682 | alpha | +/- |  |  |
| 1-2232b | beta | - |  |  |
| 661 Ulrich | beta | - | - | - |
| 00672A | beta | ++ |  |  |
| 5384 | alpha | ++ | +/- | - |
| D163 | gamma | ++ | +/- | +/- |
| Figus | double | ++ | ++ | - |
| 968A | beta | ++ |  |  |
| 131 Watkins | beta | ++ | ++ | - |
| 2698B | gamma | ++ |  |  |
| Nima | alpha | ++ | ++ | - |
| D1 | gamma | ++ | ++ | - |
| ATCC 6911 | gamma | - |  |  |

<sup>a/</sup>Following growth for 24 hrs at 35°C on 5% sheep blood agar.
$\alpha$ - hemolysis = incomplete clearing with a greenish tinge;
$\beta$ - hemolysis = complete clearing;
$\gamma$ - hemolysis = no hemolysis;
double hemolysis = inner $\beta$ - hemolysis zone with an outer $\alpha$ - hemolysis zone.

The osmotic/salt tolerance of the respective strains was tested by their ability to grow on mannitol-salt agar plates containing 7.5% NaCl. Twelve of the strains grew well, if slowly, on these plates (Table 2) but once again there did not appear to be a correlation between salt tolerance and insect pathogenicity. The three strains which could not grow on mannitol salt plates at 25°C all exhibited high group 2 pathogenicity. There was also a strong temperature effect for salt tolerance. For the 12 strains which grew well on mannitol-salt agar at 25°C, only two still grew at 37°C, and those two grew poorly (Table 2). These results are consistent with those expected for a population of *S. marcescens* [32] because all strains of *S. marcescens* ferment mannitol as the sole carbon and energy source and, even though 0.5% NaCl is the optimal salt concentration for growth, > 90% of strains grew in 7% NaCl, 11– 89% grew in 8.5% NaCl, and none grew in 10% NaCl [32]. Interestingly, the three red pigmented strains (KWN, Nima, and D1) were colorless when growing on the high salt mannitol plates at 25°C (Table 2). It is well known that pigmentation is determined in part by cultural conditions, including amino acids, carbohydrates, pH, temperatue, and inorganic ions, and that prodigiosin is not made anaerobically [32]. Now we know that prodigiosin is not made on mannitol-salt agar plates.

### Carbon and Nitrogen sources

Trehalose is the carbohydrate commonly found in insect hemolymphs. All of the strains were able to use trehalose in place of glucose and all grew well on sorbitol MacConkey (SMAC) agar plates, producing the pinkish purple colonies expected for bacteria able to utilize sorbitol [35]. One strain (5384) caused the bile salts in the SMAC plates to precipitate. The uniform ability to metabolize sorbitol was expected because *S. marcescens* is characterized by the ability to ferment sorbitol but not arabinose, rhamnose, or xylose [32, 35]. Additionally, the ability to metabolize other more unusual carbon sources helps define an organism’s biotype. Biotyping is an important tool in the epidemiology of nosocomial *S. marcescens* [24, 35] and at least 19 biotypes are now recognized [24, 32]. Our 15 *S. marcescens* strains exhibited 8 biotypes (Table 1) based on their ability to use erythritol, D, L-carnitine, benzoic acid, 3-hydroxybenzoic acid, 4-hydroxybenzoic acid, and trigonelline as their sole source of carbon and energy (S2 Table). For comparison, the 23 strains of *S. marcescens* isolated from diseased honey bee larvae in the Sudan were all biotype A4b and all unpigmented [9].

The 15 strains of *S. marcescens* were identical with regard to their nitrogen requirements. They were all able to use ammonium, nitrate, urea, uric acid, and allantoin as their sole nitrogen source, although strain 5384 used allantoin poorly. The latter 3 nitrogen sources were chosen because they could be relevant for successful growth in the insect hemolymph [36]. Uric acid is a common nitrogenous excretory product for terrestrial insects and it is often used as a nitrogen reserve [36] when insects are grown on high nitrogen diets such as those used for tobacco hornworm larvae [14]. Urea and allantoin are commonly the first products made in the degradation of uric acid.

### Extracellular Enzymes

*Serratia* sp. are well known for the production of extracellular enzymes, including chitinases [20, 37], nucleases [22], proteases [38], lipases [39], and phospholipases [39]. As determined by the size of their zones of hydrolysis on agar plates, all 15 strains of *S. marcescens* excreted DNases, proteases, lipases, and phospholipases. Proteases were detected on milk-, gelatin-, and hemoglobin-containing agar plates (S3 Table) while lipases were detected on plates containing Tweens 20, 40, 60, or 80 (S4 Table). Additionally, for the proteases, lipases, and phospholipases, one set of plates was incubated in air while a duplicate set was incubataed in a candle jar. The candle jar was chosen to simulate the high CO_2_, microaerophilic environment of insect hemolymph [40]. However, only minimal differences were observed (S3 and S4 Tables). The zones of clearing/hydrolysis for the 14 pathogenic strains of *S. marcescens* generally agreed within ± 10% while the zones for the non-pathogenic D1 were often smaller because, being non-motile, their colonies were smaller. These results are consistent with the impressive exoenzyme repertoire expected for all species of *Serratia* [32]. Our strains were also tested for chitinase production on plates containing solubilized chitin [20]. Major differences were observed (Table 1 and S4 Table), but it was the highly toxic strains which often did not have zones of clearing. In particular, strains KWN and 9674 did not excrete chitinase (Table 1).

### Autoinducers and Quorum Sensing Molecules

We next examined our collection of *S. marcescens* strains with regard to their production of acylhomoserine lactone (HSL) autoinducers. N-acyl homoserine lactone-based quorum sensing commonly regulates surfactant production [33], swarming [34, 39, 41], adhesion and biofilm formation [42] as well as the release of exoenzymes [39] and exopolysaccharides [42]. In addition, based on the precedent of other Gram negative bacteria, it could regulate an as yet unidentified insecticidal toxin. We used the thin layer chromatography overlay method described by Shaw et al [13] to identify HSLs based on color production by three reporter strains, *Agrobacterium tumefaciens, Chromobacterium violaceum*, and a strain of *Escherichia coli* responsive to the C_12_ 3-oxo HSL autoinducer made by *Pseudomonas aeruginosa*. Each of the reporter strains can respond to an exogenous autoinducer but does not produce its own autoinducer. The *Chromobacterium* reporter responds best to the C_4_, C_6_, and C_8_ 3-unsubstituted HSLs while the *Agrobacterium* reporter responds best to the C_6_, C_8_, and C_10_ 3-oxo or 3-hydroxy HSLs [13]. Our results are shown in Table 3. There were marked differences among the strains. Eight moderately insecticidal strains of clinical origin gave 1 or 2 purple spots with the *C. violaceum* detection system (Table 3). Based on their R_f_ values these spots are likely due to C_4_ and C_6_ 3-unsubstituted molecules, butanoyl and hexanoyl HSL, identified by previous researchers [39, 41, 42]. However, the highly insecticidal strains KWN and 9674 did not give any spots with the *C. violaceum* detection system. Instead, they gave a single spot with the *A. tumefaciens* detection system which is likely the 3-oxo or 3-hydroxyl C_8_ HSL while strains 968A and 2698B gave spots which are likely the 3-oxo or 3-hydroxyl C_10_ molecules (Table 3). None of the *S. marcescens* strains tested produced any molecules which reacted with the 3-oxo C_12_ specific reporter. These HSL identifications are based on comparison of their R_f_ values with known compounds and must still be considered as tentative.

**Table 3.** Autoinducer production by strains of *Serratia marcescens*.

| Strains of<br>*S. marcescens* | Pathogenicity<br>group | *C. violaceum* |  | *A. tumefaciens* |  | *E. coli* pKDT17 |
| --- | --- | --- | --- | --- | --- | --- |
|  |  | #AI | R<sub>f</sub> | #AI | R<sub>f</sub> | #AI |
| KWN | 1 | 0<sup>c</sup> |  | 1<sup>b</sup> | 0.39 | 0 |
| 9674 | 1 | 0<sup>c</sup> |  | 1<sup>b</sup> | 0.32 | 0 |
| E223 | 2 | 0 |  | ND | ND | ND |
| 1682 | 2 | 1 | 0.45 | 0 | 0 | 0 |
| 1-2232b | 2 | 2 | 0.44 & 0.67 | ND | ND | ND |
| 661 Ulrich | 2 | 2 | 0.42 & 0.62 | ND | ND | ND |
| 00672A | 3 | 2 | 0.46 & 0.66 | ND | ND | ND |
| 5384 | 3 | 2 | 0.47 & 0.72 | ND | ND | ND |
| D163 | 3 | 2<sup>d</sup> | 0.57 & 0.68 | ND | ND | 0 |
| Figus | 3 | 1 | 0.50 | ND | ND | ND |
| 968A | 4 | 2 | 0.44 & 0.66 | 1 | 0.15 | 0 |
| 131 Watkins | 4 | 0 |  | ND |  | ND |
| 2698B | 5 | 0 |  | 2 | 0.11 & 0.48 | 0 |
| NIMA | 5 | 0 |  | ND |  | ND |
| D1 | 6 | 0 |  | ND |  | ND |
| Positive Controls |  |  |  |  |  |  |
| *C. violaceum* |  | 3 |  | 0 |  | ND |
| *P. aeruginosa* (PA01) |  | 1 |  | 1 |  | 1 |
| *Vibrio fischeri* |  | 1 or 2 |  | ND |  | ND |

A/ Tentative identities were based on the R<sub>f</sub> values reported by Shaw et al [13] of 0.82, 0.68, 0.41, 0.18, and 0.07 for the C<sub>4</sub>, C<sub>6</sub>, C<sub>8</sub>, C<sub>10</sub>, and C<sub>12</sub> 3-oxo HSLs, respectively, and 0.77, 0.47, 0.23, 0.09, and 0.02 for the C<sub>4</sub>, C<sub>6</sub>, C<sub>8</sub>, C<sub>10</sub>, and C<sub>12</sub> 3-unsubstituted HSLs, respectively. ND = not determined. Note that the R<sub>f</sub> values reflect migration on the C<sub>18</sub> reversed phase TLC plates and thus should apply equally for all three detection systems.
B/ For cells grown on trypticase-soy broth, none detected for cells grown on minimal ccy medium [25].
C/ None detected for cells grown either on trypticase-soy broth, or on minimal ccy medium [25].
D/ For cells grown on minimal ccy medium, only one AI (R<sub>f</sub> = 0.46) was detected.

### Antibiotic Sensitivity/Resistance

Antibiotic resistance profiles were determined for 16 strains of *S. marcescens* using 24 antibacterials for which antibiotic discs were commercially available (Table 4). The profile for the quality control strain *E. coli* 23744 was as expected (Table 4). The 15 strains for which we had determined insect pathogenicity were resistant to 12-17 of the 24 antibiotics (R + NZ). The strains were uniformly resistant to tobramycin, tetracycline, chloramphenicol, erythromycin, rifampin, nitrofurantoin, novobiocin, ampicillin, bacitracin, vancomycin, polymyxin B, and colistin (Table 4). The uniform resistance to erythromycin, novobiocin, bacitracin, and vancomycin is expected because these antibiotics are usually only active against Gram positive bacteria. The uniform resistance to ampicillin, colistin, tetracycline,and polymyxin B is characteristic of most Serratia [32, 35]. The only antibiotics for which insect pathogenicity and antibiotic resistance correlated were cefoxitin and cefotaxime, where the pathogenic strains (groups 1-4) were resistant while the non-pathogenic groups 5-6 were not, Finally, the results for sulfamethoxazole/trimethoprim are intriguing in that they alternate between sensitivity (S/I) and complete resistance (NZ) in a seemingly random fashion (Table 4). However, they do not correlate with either insect pathogenicity or biotype and, consequently, we have not investigated them further.

**Table 4.** Antibiotic resistance/sensitivity of *Serratia marcescens*.

| Strain | Protein synthesis |  |  |  |  |  |  |  | Nucleic acid |  |  |  |  |  | Cell wall |  |  |  |  |  |  |  | Membrane |  | Folic acid |
| --- | --- | --- | --- | --- | --- | --- | --- | --- | --- | --- | --- | --- | --- | --- | --- | --- | --- | --- | --- | --- | --- | --- | --- | --- | --- |
|  |  | S | K | N | NN | TE | C | E | RA | NA | CIP | NOR | F/M | NB | AM | PIP | FOX | CTX | ATM | IPM | B | VA | PB | CL | SXT |
| E. coli 23744 | - | I | I | R | R | S | S | NZ | - | S | S | S | S | NZ | S | S | S | S | S | S | R | I | I | NZ | S |
| KWN | 1 | R | I | R | NZ | I | R | NZ | R | S | S | S | NZ | NZ | R | S | R | I | S | S | NZ | R | R | NZ | I |
| 9674 | 1 | I | I | R | NZ | R | NZ | NZ | R | S | S | S | NZ | R | R | S | R | R | S | S | NZ | R | NZ | NZ | NZ |
| E223 | 2 | I | I | I | R | NZ | R | NZ | R | S | S | S | NZ | NZ | NZ | I | R | R | S | S | NZ | NZ | NZ | NZ | NZ |
| 1682 | 2 | I | I | I | R | R | R | NZ | R | S | - | S | NZ | NZ | NZ | I | I | S | S | S | NZ | R | NZ | NZ | I |
| 1-2232b | 2 | NZ | R | R | NZ | - | R | NZ | R | S | S | S | NZ | NZ | R | S | R | NZ | S | S | NZ | NZ | NZ | NZ | NZ |
| 661 Uhlich | 2 | R | - | - | NZ | R | I | NZ | R | S | S | S | NZ | R | NZ | R | R | NZ | S | S | NZ | R | NZ | NZ | I |
| 00672A | 3 | I | NZ | NZ | R | NZ | NZ | NZ | R | S | S | S | NZ | NZ | NZ | S | R | NZ | S | S | NZ | NZ | NZ | NZ | NZ |
| 5384 | 3 | - | R | R | R | I | NZ | NZ | R | S | S | S | NZ | NZ | R | S | R | R | S | S | NZ | R | NZ | NZ | NZ |
| D163 | 3 | I | I | R | R | NZ | R | NZ | R | S | S | S | NZ | NZ | R | S | R | I | S | S | NZ | NZ | NZ | NZ | NZ |
| Figus | 3 | R | I | R | NZ | R | R | NZ | R | S | S | S | NZ | NZ | R | S | R | R | S | S | NZ | R | NZ | NZ | I |
| 698A | 4 | I | R | R | NZ | R | - | NZ | R | S | I | S | NZ | NZ | NZ | S | R | R | S | S | NZ | R | NZ | NZ | I |
| 131 Watkins | 4 | I | I | I | NZ | R | R | NZ | R | S | S | S | NZ | NZ | NZ | S | R | NZ | S | S | NZ | R | NZ | NZ | NZ |
| 2698B | 5 | I | I | R | R | R | NZ | NZ | R | S | S | S | NZ | NZ | S | S | I | - | S | S | NZ | NZ | NZ | NZ | I |
| Nima | 5 | I | I | R | NZ | R | NZ | NZ | R | S | S | S | NZ | NZ | R | S | I | I | S | S | NZ | R | R | NZ | S |
| D1 | 6 | NZ | - | R | R | R | R | NZ | R | S | S | S | NZ | NZ | I | S | S | S | S | S | NZ | R | NZ | NZ | S |
| ATCC 6911 | - | I | I | R | NZ | I | I | NZ | R | S | S | S | NZ | NZ | NZ | S | S | S | S | S | NZ | R | NZ | NZ | S |
| Resistant |  | <11 | <13 | <12 | <12 | <14 | <12 | <13 | <16 | <13 | <15 | <12 | <14 | <17 | <13 | <17 | <14 | <14 | <8 | <13 | <8 | <14 | <8 | <18 | ≤10 |
| Sensitive |  | >15 | >18 | >17 | >15 | >19 | >18 | >23 | >20 | >19 | >21 | >17 | >17 | >17 | >17 | >21 | >18 | >18 | >23 | >16 | >13 | >17 | >12 | >11 | >16 |

a/ S=streptomycin; K=kanamycin; N=neomycin; NN=tobramycin; TE=tetracycline; C=chloramphenicol; E=erythromycin; RA=rifampin; NA=nalidixic acid; CIP=ciprofloxacin; NOR=norfloxacin; F/M=nitrofurantoin; NB=novobiocin; AM=ampicillin; PIP=piperacillin; FOX=cefoxitin; CTX=cefotaxime; ATM=aztreonam; IPM=imipenem; B=bacitracin; VA=vancomycin; PB=polymyxin B; CL=colistin; SXT=sulfamethoxazole/trimethoprim.
b/ S=sensitive; I=intermediate; R=resistant; NZ=no zone of inhibition at all; - = not done.
c/ Column 2 = Insect Pathogenicity Groups

#### Bacteriophage Sensitivity/Resistance

Each strain of *S. marcescens* was tested versus a collection of seven broad host range bacteriophage (SN-1, SN-2, SN-X, SN-T, BHR1, BHR2, and D_3_ C_3_), known to be active versus multiple Gram negative bacteria [28]. The SN and BHR bacteriophage had originally been isolated from *Sphaerotilus natans* and *Pseudomonas aeruginosa*, respectively [28]. We previously showed [28] that these broad host range bacteriophage, as produced on either *P. aeruginosa* or *S. natans*, were unable to infect *S. marcescens* KWN [28]. We now observed that they were also unable to infect or propagate on any of the 15 strains listed in Table 1. No plaques were produced on agar plates and no increases in phage titer were observed in liquid culture (data not shown). These findings are consistent with the generalization of Grimont and Grimont [32] that bacteriophages isolated on genera other than *Serratia* rarely multiply on *Serratia*. These phage sensitivity screens were conducted in part in the hope of identifying effective biocontrol mechanisms for *S. marcescens* but also as an indirect method for determining the presence and importance of Type IV pili in insect pathogenicity. The Type IV secretion system is a well known virulence mechanism allowing bacteria to inject protein toxins into other cell types while phage SN-T was shown to be broad host range because it attached to various bacteria by means of their Type IV pili [43]. Thus, the absence of phage infectivity provides no evidence for Type IV pili under laboratory conditions but does not rule out a role for Type IV secretion in insect pathogenicity.

## Discussion

Fifteen strains of *S. marcescens* were examined for their insect pathogenicity towards *M. sexta* larvae. They fell into six groups, ranging from the highly toxic Group 1 to non-toxic Group 6 (Table 1). The need for comparatively high inocula (5 × 10^5^ bacteria per larva) to achieve these variable LD_50_ values reflects both the large size of the larvae and their strong antibacterial defense mechanisms, worthy of study on the methods employed by the strains of *S. marcescens* to overcome them. We presume that the virulence gradient exhibited by these 15 strains of *S. marcescens* is not specific for *M. sexta* but instead extends to other insects as well. For instance, preliminary evidence (Brian Lazzaro, Cornell, personal communication) confirms that the *S. marcescens* strains toxic to *M. sexta* (Groups 1-4) were also virulent to adult *D. melanogaster* while those not virulent to *M. sexta* (Groups 5-6) were not virulent to *D. melanogaster*. The virulence properties of these 15 strains are also being tested versus the red flour beetle *Tribolium castaneum* (Ann Tate, Vanderbilt, personal communication).

Relative insect toxicity was not correlated with pigmentation, colony morphology, biotype, motility, capsule formation, iron availability, surfactant production, swarming ability, antibiotic resistance, bacteriophage susceptibility, salt tolerance, nitrogen utilitization patterns, or the production of 4 exoenzymes: proteases, DNase, lipase, or phospholipase. Thus, we found no relationship between traditional pathogenicity factors and observed insect pathogenicity. However, the absence of motility in strain D1 probably explains in part its non-pathogenic status. In this regard, it is important that we tested pathogenicity by injecting the bacterial cells directly into the insect hemolymph. Under these conditions, four of the six highly pathogenic (Groups 1 and 2) strains were chitinase negative (Table 1). The relative importance of extracellular chitinase would likely have been different if the bacterial cells had been provided in the diet or sprayed on the larval surface where they would have had to penetrate the peritrophic membrane or the cuticle to exert their pathogenicity. We note that Ruiz-Sanchez et al [44] compared 102 strains of *S. marcescens* for their chitinolytic activity. They found that *S. marcescens* Nima, which exhibited very little pathogenicity in our assays (Fig. 1), had ca. 43 times higher chitinolytic activity than most other *S. marcescens* strains [44].

Pathogenicity did, however, correlate with the type of autoinducer produced (Table 3). *S. marcescens*, like a great many Gram negative bacteria [45], is known to produce HSL autoinducers which act in a quorum sensing manner [41, 46]. Bainton et al [46] used an autoinducer-dependent bioluminescence system to detect 3-oxo-hexanoyl HSL activity in *S. marcescens* supernatants. The identity of this molecule was confirmed by infrared, mass spectrometric, and NMR analysis [46]. Later, Eberl et al [41] showed that *Serratia liquefaciens*, now *S. marcescens* [42], produced butanoyl and hexanoyl HSLs in a ratio of 10:1 and these autoinducers controlled the differentiation to swarming motility. In this case, the autoinducers without a side chain oxygen at position 3 were more effective in causing swarming than those with a 3-oxo side chain [41]. The distinction is relevant because: A/ the strain of *S. liquefaciens* used [41] has been reclassified as *S. marcescens* based on its 16S rRNA sequence [42]; and B/ the *Agrobacterium* based TLC detection system strongly prefers the 3-oxo homoserine lactones whereas the *Chromobacterium* based system strongly prefers the 3-unsubstituted molecules [13]. Based on a comparison with R_f_ values of known HSLs [13], the hightly insecticidal strains KWN and 9674 (Group 1) produced 3-oxo or 3-hydroxy C_8_ HSL while the moderately insecticidal clinical isolates, 1682 through 968A in Table 3, produced one or both of the 3-unsubstituted C_4_ and C_6_ HSLs (Table 3), the same HSLs as found by Eberl et al [41]. Finally, the poorly insecticidal strains 968A and 2698B produced 3-oxo hydroxy C_10_ HSL. We believe that the 3-oxo or hydroxy C_8_ and C_10_ HSLs are autoinducers not previously reported from *Serratia* sp.

Unfortunately we have as yet little evidence regarding which genes or virulence factors are being regulated by the respective autoinducers. The extracellular nuclease of *S. marcescens* is regulated in a growth-phase and cell-density dependent manner [22] as are all the exoenzymes produced by *S. liquefaciens* [39], now *S. marcescens* [42], and the C_4_ and C_6_ HSL autoinducers also regulate surfactants [19, 33], swarming [34, 39, 41], adhesion, biofilms, and exopolysaccharide production [42]. Some strains of *S. marcescens* also produce a heat-labile enterotoxin as a virulence factor [47] which could also be regulated in a quorum sensing manner. An additional complication is that all of our studies were conducted with typical bacterial growth media under aerobic growth conditions. However, since the HSLs produced can vary depending on the growth media and growth conditions [13 and Table 3], we have no assurance the HSLs detected are the same ones which would be produced in the insect gut or hemolymph.

Finally, many insects including *M. sexta* produce antimicrobial peptides (AMPs) which are essential components of insect immunity [48, 49]. These peptides are typically low molecular weight, heat stable, cationic, and produced in the fat body of insects. They insert into and disrupt microbial membranes, thereby promoting pathogen clearance and insect survival [48]. AMPs have been studied in Drosophila melanogaster [49], *M. sexta* [50, 51], and many other insects. Significantly, these AMPs are localized in the larval hemolymph and in many cases mutants unable to produce those AMPs have become susceptible to microbial infection [49]. Thus, a logical series of experiments would be to compare the susceptibilities of these 15 strains of *S. marcescens* to the individual AMPs produced by *M. sexta* [50, 51] with the expectation that the highly pathogenic strains would be either more resistant to those AMPs or somehow avoid induction of AMP expression by the larvae. It is our hope that the molecular features distinguishing our 15 strains of *S. marcescens* will prove useful in combatting insect diseases such as colony collapse disorder of honey bees [10].

## Acknowledgements

We thank Lynda Liska for her advice on the care and rearing of *Manduca sexta*, and Daniel Nickerson for preparing Fig 1.

## Supporting information

**Table S1.**
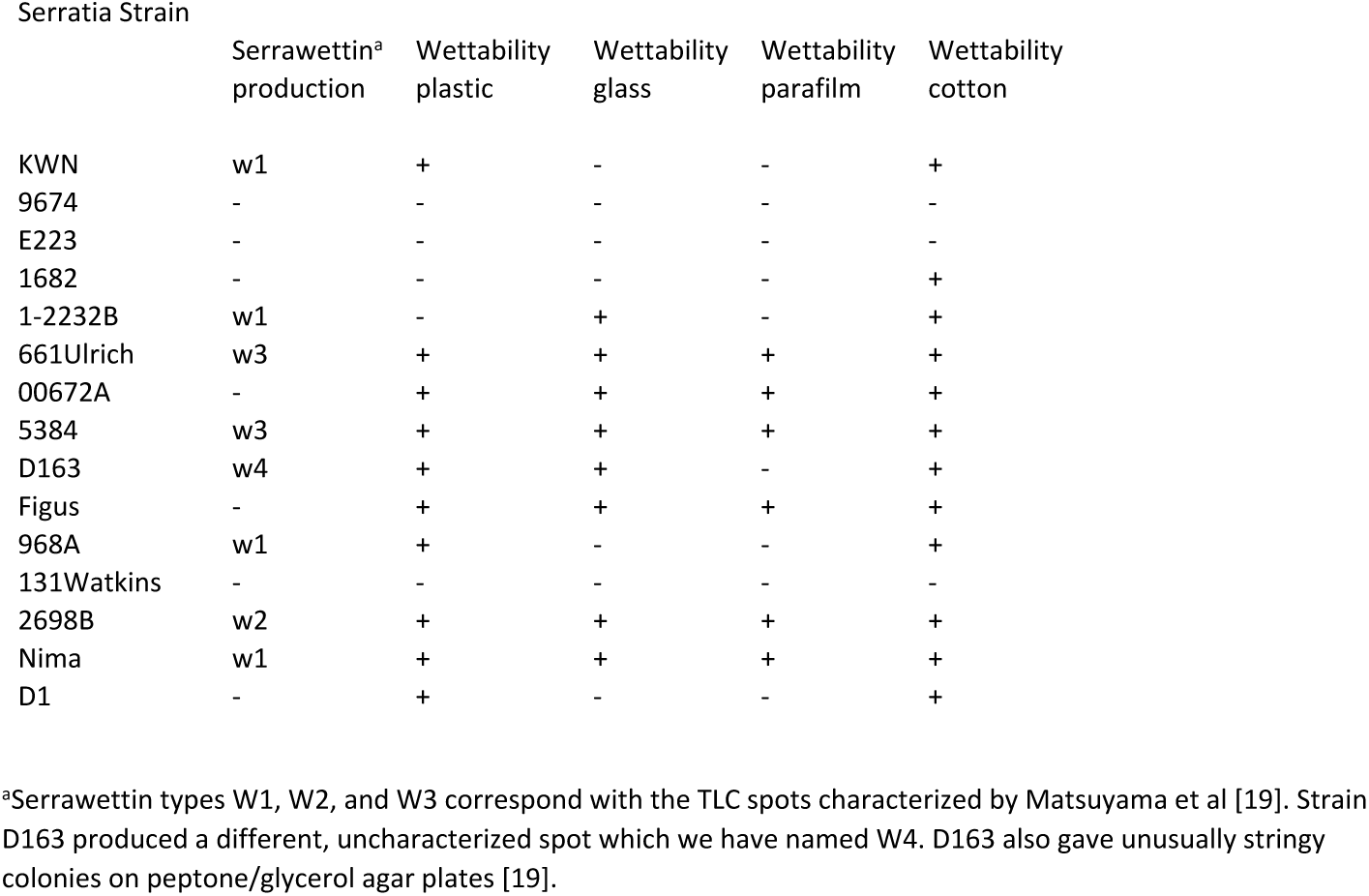
Surfactants and wettability.

| Serratia Strain | Serrawettin<sup>a</sup><br>production | Wettability<br>plastic | Wettability<br>glass | Wettability<br>parafilm | Wettability<br>cotton |
| --- | --- | --- | --- | --- | --- |
| KWN | w1 | + | - | - | + |
| 9674 | - | - | - | - | - |
| E223 | - | - | - | - | - |
| 1682 | - | - | - | - | + |
| 1-2232B | w1 | - | + | - | + |
| 661Ulrich | w3 | + | + | + | + |
| 00672A | - | + | + | + | + |
| 5384 | w3 | + | + | + | + |
| D163 | w4 | + | + | - | + |
| Figus | - | + | + | + | + |
| 968A | w1 | + | - | - | + |
| 131Watkins | - | - | - | - | - |
| 2698B | w2 | + | + | + | + |
| Nima | w1 | + | + | + | + |
| D1 | - | + | - | - | + |

<sup>a</sup>Serrawettin types W1, W2, and W3 correspond with the TLC spots characterized by Matsuyama et al [19]. Strain D163 produced a different, uncharacterized spot which we have named W4. D163 also gave unusually stringy colonies on peptone/glycerol agar plates [19].

**Table S2.** Carbon Sources used to determine biotypes.

| Serratia Strain |  |  |  |  | 4-OH<br>benzoic<br>acid | 3-OH<br>benzoic<br>acid | Prodigiosin | Bio type |
| --- | --- | --- | --- | --- | --- | --- | --- | --- |
|  | Erythritol | Benzoic<br>acid | Carnitine | Trigonelline |  |  |  |  |
| KWN | + | + | + | - | - | - | + | A1a |
| 9674 | - | - | - | + | + | + | - | A8b |
| E223 | - | - | + | + | - | - | - | TCT |
| 1682 | - | - | + | + | + | - | - | A5 |
| 1-2232B | + | - | + | - | + | - | - | A4a |
| 661Ulrich | + | - | + | - | + | - | - | A4a |
| 00672A | + | - | + | - | + | - | - | A4a |
| 5384 | - | - | - | + | + | - | - | A8a |
| D163 | + | - | + | - | + | - | - | A4a |
| Figus | + | - | + | - | + | - | - | A4a |
| 968A | + | - | + | - | + | - | - | A4a |
| 131Watkins | - | - | - | + | + | - | - | A8a |
| 2698B | + | - | + | + | - | + | - | A3b |
| Nima | + | - | + | - | - | - | + | A2a |
| D1 | + | - | - | - | - | - | + | A2a |

**Table S3.** Protease and Phosphatidyl Choline Assays.

| Serratia Strain | phosphatidylcholine |  | gelatin |  | milk |  | hemoglobin |  |
| --- | --- | --- | --- | --- | --- | --- | --- | --- |
|  | air | candle jar | air | candle jar | air | candle jar | air | candle jar |
| KWN | 22 | 21 | 12 | 15 | 9 | 9 | - | + |
| 9674 | 18 | 16 | 10 | 10.5 | 6.5 | 5 | - | + |
| E223 | 19 | 22 | 13 | 13 | 8.5 | 6 | - | + |
| 1682 | 19 | 15 | 9 | 12.5 | 6 | 7 | - | + |
| 1-2232B | 22 | 21.5 | 11.5 | 11 | 7 | 7 | + | + |
| 661Ulrich | 21 | 20 | 10 | 11 | 6.5 | 6 | - | + |
| 00672A | 23 | >22 | 12 | 12.5 | 7.5 | 9 | + | + |
| 5384 | 14.5 | 9 | 12 | 11.5 | 8 | 6 | - | + |
| D163 | 22.5 | 23 | 10 | 11 | 7 | 7 | + | + |
| Figus | 22 | + | 12 | 13 | 8 | 7 | + | + |
| 968A | 23 | 22.5 | 11.5 | 14 | 9 | 7.5 | + | - |
| 131Watkins | 17.5 | 17.5 | 12 | 12 | 8 | 8 | - | - |
| 2698B | 25 | 21 | 10 | 12 | 8 | 6.5 | - | + |
| Nima | 16 | + | 10 | 11 | 5 | 8.5 | + | - |
| D1 | 12 | 12 | 9 | 11 | 8 | 6 | + | + |

**Table S4.** Chitinase and Lipase Assays.

| Serratia Strain | Motility | Chitinase<br>production<br>yes/no | Chitinase<br>production<br>zone/dia. | Lipase:<br>Tween 20<br>air | Lipase:<br>Tween 20<br>candle jar | Lipase:<br>Tween 40<br>air | Lipase:<br>Tween 40<br>candle jar | Lipase:<br>Tween 60<br>air | Lipase:<br>Tween 60<br>candle jar | Lipase:<br>Tween 80<br>air | Lipase:<br>Tween 80<br>candle jar |
| --- | --- | --- | --- | --- | --- | --- | --- | --- | --- | --- | --- |
| KWN | + | - |  | 30.5 | 23 | 29.75 | 29 | 27.5 | 30 | 28 | 22.5 |
| 9674 | + | - |  | 27.5 | 22 | 28.5 | 27 | 27.75 | 24 | 28.5 | 28 |
| E223 | + | + | 10mm | 26.75 | 20 | 32.5 | 33 | 28.5 | 28 | 27 | 23 |
| 1682 | + | + | 8mm | 26.25 | 19 | 28.5 | 28 | 26.5 | 21 | 25 | 21 |
| 1-2232B | + | - |  | 30.5 | 29 | 34.5 | 33.5 | 27 | 32 | 31 | 25 |
| 661Ulrich | + | - |  | 29.75 | 28.5 | 31 | 31 | 30 | 24 | 28.5 | 28 |
| 00672A | + | + | 7.5mm | 31 | 30 | 33.25 | 34 | 31 | 29 | 30.5 | 24 |
| 5384 | + | + | 8.5mm | 27.25 | 17.5 | 29 | 31 | 29 | 24 | 25.75 | 0 |
| D163 | + | + | 10mm | 30.75 | 23.5 | 30.5 | 34 | 32.75 | 26 | 31.5 | 25.5 |
| Figus | + | + | 8mm | 29 | 26 | 33.5 | 39 | 25.75 | 28 | 30.5 | 24.5 |
| 968A | + | + | 6mm | 31 | 23.5 | 34.5 | 32.5 | 30.5 | 25 | 29.5 | 25 |
| 131Watkins | + | + | 12mm | 29 | 26.5 | 28.5 | 33 | 28.5 | 23 | 25.5 | 25 |
| 2698B | + | + | 15.5mm | 31.75 | 28 | 35 | 28 | 29.5 | 29 | 29 | 30 |
| Nima | + | + | 13mm | 30.5 | 23 | 27.5 | 27.5 | 29.5 | 24 | 29.5 | 23 |
| D1 | - | + | 7.5mm | 20.5 | 18.5 | 17 | 23 | 24.5 | 20 | 19.5 | 20 |

